# A toolbox of immunoprecipitation-grade monoclonal antibodies against human transcription factors

**DOI:** 10.1101/116442

**Authors:** Anand Venkataraman, Kun Yang, Jose Irizarry, Mark Mackiewicz, Paolo Mita, Zheng Kuang, Lin Xue, Devlina Ghosh, Shuang Liu, Pedro Ramos, Shaohui Hu, Diane Bayron, Sarah Keegan, Richard Saul, Simona Colantonio, Hongyan Zhang, Florencia Pauli Behn, Guang Song, Edisa Albino, Lillyann Asencio, Leonardo Ramos, Luvir Lugo, Gloriner Morell, Javier Rivera, Kimberly Ruiz, Ruth Almodovar, Luis Nazario, Keven Murphy, Ivan Vargas, Zully Ann Rivera-Pacheco, Christian Rosa, Moises Vargas, Jessica McDade, Brian S. Clark, Sooyeon Yoo, Seva G. Khambadkone, Jimmy de Melo, Milanka Stevanovic, Lizhi Jiang, Yana Li, Wendy Y. Yap, Brittany Jones, Atul Tandon, Elliot Campbell, Stephen Anderson, Richard M. Myers, Jef D. Boeke, David Fenyo, Gordon Whiteley, Joel S. Bader, Ignacio Pino, Daniel J. Eichinger, Heng Zhu, Seth Blackshaw

## Abstract

A key component to overcoming the reproducibility crisis in biomedical research is the development of readily available, rigorously validated and renewable protein affinity reagents. As part of the NIH Protein Capture Reagents Program (PCRP), we have generated a collection of 1406 highly validated, immunoprecipitation (IP) and/or immunoblotting (IB) grade, mouse monoclonal antibodies (mAbs) to 736 human transcription factors. We used HuProt™ human protein microarrays to identify mAbs that recognize their cognate targets with exceptional specificity. Using an integrated production and validation pipeline, we validated these mAbs in multiple experimental applications, and have distributed them to the Developmental Studies Hybridoma Bank (DSHB) and several commercial suppliers. This study allowed us to perform a meta-analysis that identified critical variables that contribute to the generation of high quality mAbs. We find that using full-length antigens for immunization, in combination with HuProt™ analysis, provides the highest overall success rates. The efficiencies built into this pipeline ensure substantial cost savings compared to current standard practices.

## Introduction

Specific affinity reagents are essential for studying protein function. Antibodies represent the vast majority of protein affinity reagents used by the research community. In recent years, it has become clear that there are serious problems with the quality, consistency, and availability of research-grade antibodies^1^. It is estimated that globally, over $800 million (US dollars) is wasted annually as a result of using poor quality antibodies in research^2^. This problem is aggravated by several factors, including the absence of standardized validation criteria in the research community, a lack of transparency from commercial antibody suppliers about their products, the extensive use of non-renewable polyclonal reagents which show extensive batch variation, and a lack of technologies that allow antibody cross-reactivity to be comprehensively assessed^3-7^.

In recognition of this problem, several recent efforts have been undertaken to generate large collections of high-quality antibodies against human proteins. Despite the successes of these initiatives, they nonetheless faced major limitations. While several efforts have generated renewable reagents -- either in the form of conventional monoclonals or recombinant antibodies -- these have targeted only a limited subset of proteins^8-11^. Other more comprehensive efforts generated non-renewable polyclonal antibodies^12^. Furthermore, animal-based approaches have generally used linear or denatured antigens for immunization, which yields reagents that are useful in immunoblotting and immunostaining, but whose ability to recognize native antigen for applications such as immunoprecipitation is uncertain^12^. Finally, despite often extensive validation, proof that these reagents do indeed exclusively recognize their intended targets is lacking.

To extend these efforts, the NIH Common Fund in 2010 initiated the Protein Capture Reagent Program (PCRP), an effort that was aimed at generating highly specific, extensively characterized and readily available renewable affinity reagents that target a broad range of human transcription factors and associated proteins^13^. In this manuscript, we describe an integrated production and validation pipeline for high-throughput and low-cost generation of mouse monoclonal antibodies (mAbs) that recognize their targets with both high specificity and affinity. By using human proteome microarray validation as an initial screen, we show that we can rapidly enrich for ultraspecific, high-affinity mAbs that perform well in downstream applications -- particularly immunoprecipitation. We demonstrate that protein microarray-based validation of antibody specificity can be used to identify mAbs that selectively recognize their intended target in either native or denatured conformation and do not substantially cross-react with other proteins. We demonstrate that this approach can be used to rapidly and efficiently identify proteins that cross-react with commercially available mAbs. Array-based analysis early in the development process can substantially reduce the costs incurred when non-specific mAbs are taken through validation steps.

We have used this pipeline to generate 1406 mAbs that target a total of 736 unique human transcription factors and transcription factor-associated proteins, and which work for one or more common research applications, including immunoprecipitation, immunoblotting and ChIP-Seq. We have made these mAbs readily available to the research community through both the Developmental Studies Hybridoma Bank (DSHB) and various commercial suppliers. These reagents serve as a standardized toolbox for biochemical analysis of transcriptional regulatory mechanisms in human cells.

## Results

### Design of the production pipeline

The workflow for production, validation and distribution of highly specific mAbs is shown in Figure 1. Following identification of target proteins, recombinant domains or full-length proteins were produced and purified from *E. coli* or S. *cerevisiae* respectively, and used for immunization of mice through either an intraperitoneal (i.p.) or footpad (f.p.) route of administration. Following screening for IgG-positive hybridomas using ELISA, supernatants are then subjected to a two-step microarray-based screen. First, hybridoma supernatants are screened against a microarray comprised of the antigen used for immunization and up to 80 other proteins --referred to here as a minichip. Antibodies that recognize their cognate antigen as the top target on the minichip are then subjected to specificity analysis using HuProt™ human protein microarrays (hereafter referred to as HuProt), which contain from 17,000-19,000 full-length, purified human proteins expressed in yeast^14^. Antibodies that pass this primary specificity analysis are then subjected to secondary screening, and are tested by immunoblotting and immunoprecipitation against their cognate target expressed at a range of different concentrations in human cell lines. A subset of passing mAbs are then tested by ChIP-Seq in ENCODE cell lines, or by IHC analysis against mouse and/or human tissue. Validation data for all mAbs is made available via the PCRP web portal (http://proteincapture.org). Finally, mAbs that work in one or more research-grade application are made available to the research community through DSHB or commercial distributors.

**Figure 1.**
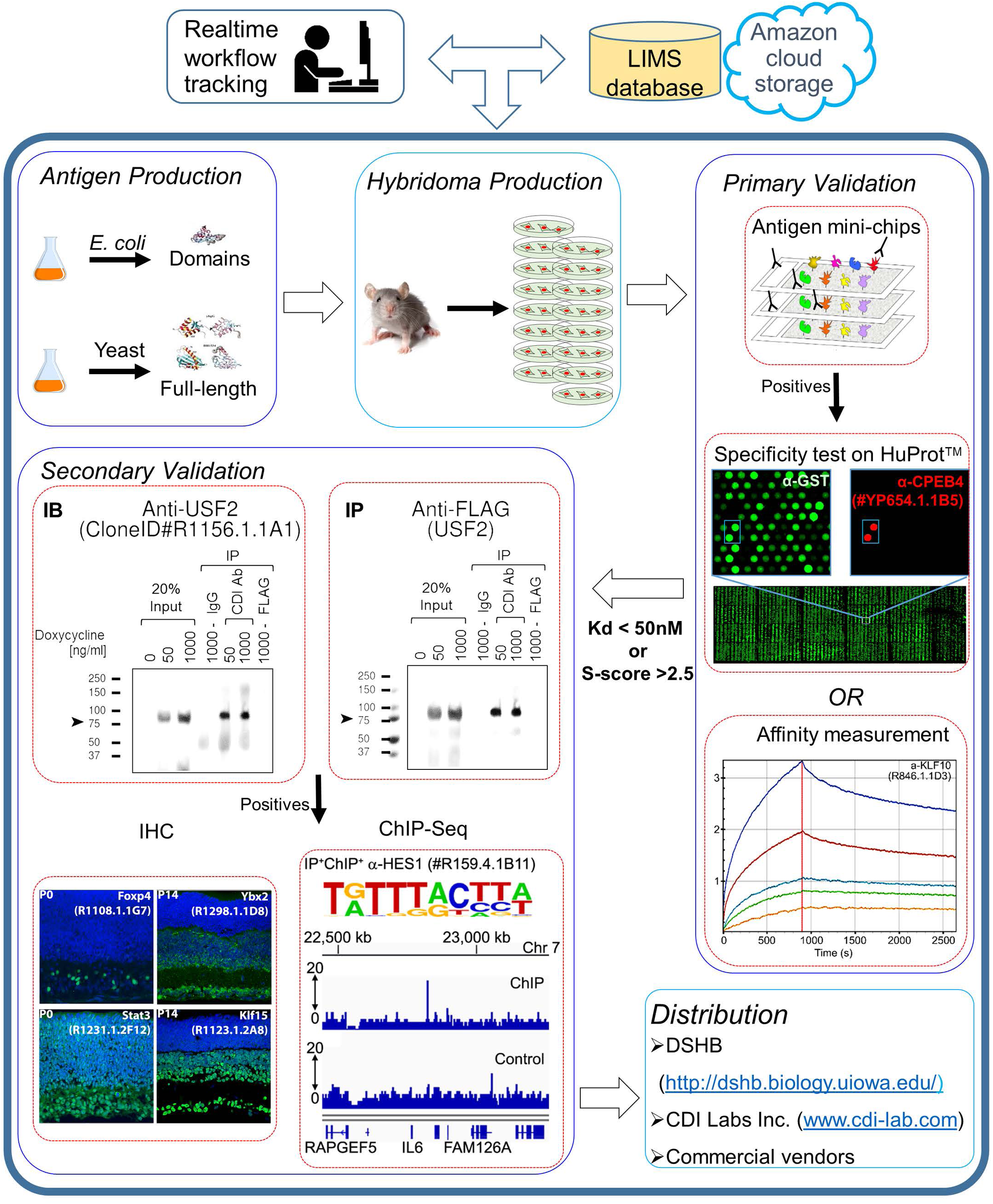
Pipeline description. A schematic overview highlighting the critical steps involved in generation and distribution of high quality monoclonal antibodies generated in this project.

### Recombinant proteins on HuProt arrays comprise a majority of the annotated, full-length proteome

By probing HuProt with individual mAbs, we not only assessed the ability for a given mAb to recognize its intended target, but at the same time counter-screened for its cross-reactivity against other proteins. To this end, it is critical that a large fraction of the human proteome is represented on the array. We performed extensive annotation and verification of our array (see online Methods). When classified based on GO annotations associated with reviewed proteins of UniProt database, >75% of proteins in each major functional category was represented on the array (Figure 2a). To the best of our knowledge, the HuProt represents the largest collection of unique proteins printed on a single array.

**Figure 2.**
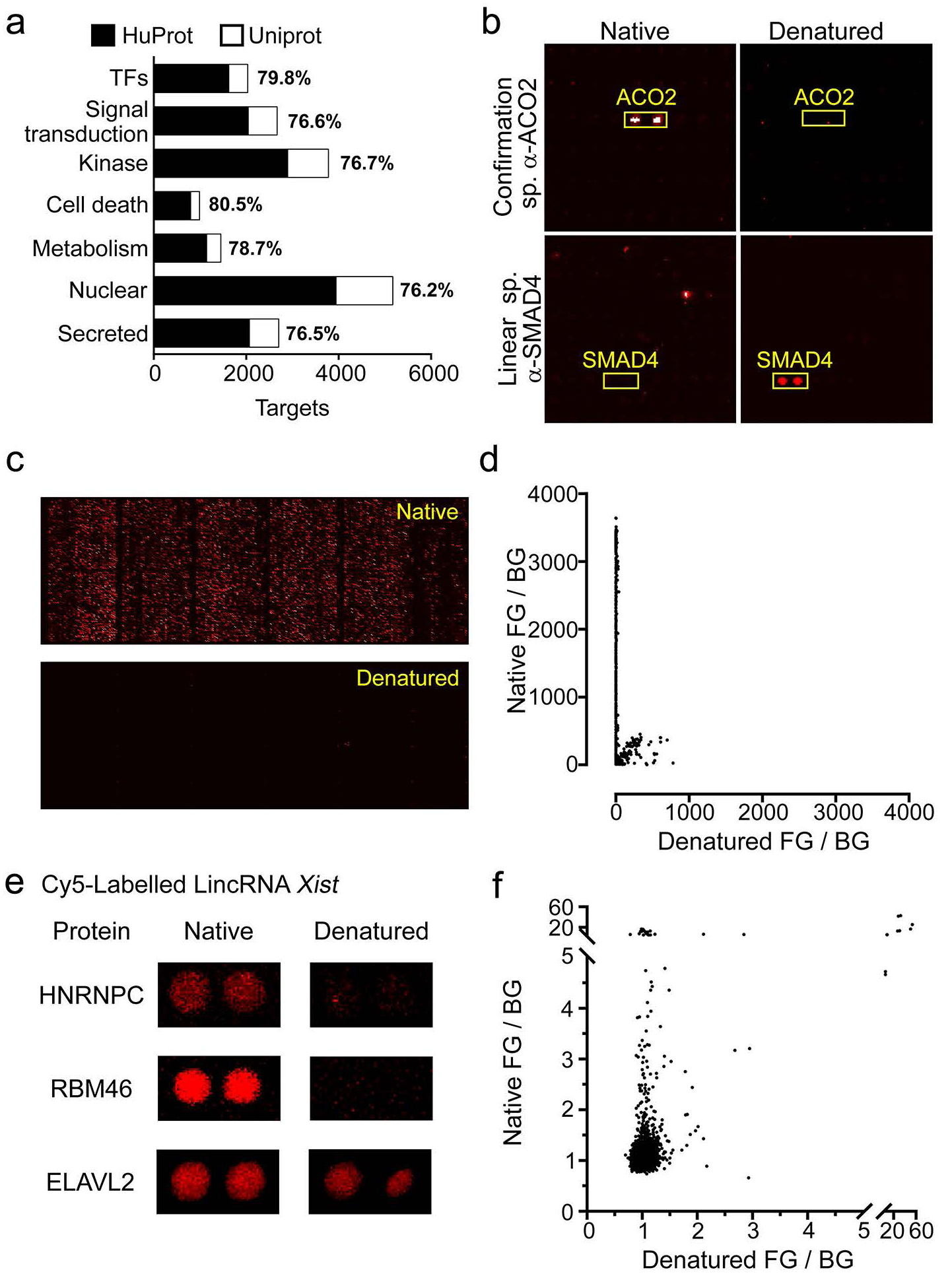
HuProt proteins are in native conformation. (a) The majority of the reviewed UniProt proteins are represented on HuProt. Categories of proteins represented are based on GO annotation. (b) Anti-ACO2 6D1BE4 mAb (Catalog #ab110320, Abcam), which recognizes a conformational specific epitope, only recognizes its cognate target on the native HuProt but not in HuProt denatured with 9M Urea and 5 mM DTT. Similarly, an anti-SMAD4 mAb (CDI Labs, #R516.2.1G11), which recognizes a linear epitope, only recognizes its cognate target on the denatured but not native HuProt. (c,d) GST-tagged proteins on the native HuProt are recognized by the conformation specific anti-GST antibody (CDI Labs, #27.3.6G8) but is almost entirely lost following denaturation of the HuProt. Plotting the signal measured from the native and denatured HuProt highlights the range of signal observed under native condition and the dramatic loss of signal following denaturation (e,f). A functional study to identify protein interaction partners of the long noncoding RNA (lincRNA) *Xist* using the native HuProt. (e) Micrographs representing the HuProt signal observed for known *Xist* interaction partner (HNRNPC) and potential new partners identified in our screen (RBM46 and ELAVL2). (f) Majority of these functional interactions are lost when *Xist* is probed against a denatured HuProt.

### Recombinant proteins on HuProt arrays are in native conformation

Since immunoprecipitation-grade mAbs need to be able to recognize their target in native conformation, we first tested whether spotted proteins used in generating the HuProt were themselves in native conformation, and therefore useful for screening mAbs. This was tested using a variety of approaches.

First, we probed the array with antibodies that had previously been shown to selectively recognize either linear or folded epitopes on their cognate antigen. We tested these antibodies on both native arrays and arrays denatured by treatment with 8 M urea and 5 mM DTT. For example, we observe that a mAb that specifically recognizes a linear epitope on SMAD4 only recognizes its target on denatured arrays, while a mAb that specifically recognizes folded ACO2 recognizes its target only on the native array (Figure 2b). In the course of our work, we identified a conformation-specific mAb to glutathione-S-transferase (GST), which preferentially recognizes the GST tag fused to the N-terminus of all targets on native arrays (Figure 2c) and shows almost no signal on the denatured array (Figure 2d), supporting the assertion that proteins are indeed in native conformation on the HuProt array. From this point on, we used this mAb to confirm that batches of HuProt arrays used for analysis were indeed in native conformation.

Finally, to determine that proteins on the array retained biochemical function, we used a sub-array containing a subset of the HuProt proteins and performed analysis to identify proteins that selectively bound the long noncoding RNA *Xist.* Using previously described methods (see online Methods), we identified 247 proteins as potential interactors with *Xist* RNA in the native HuProt array with with z-score>2 (Table S1). We identified 5 proteins that are well defined as *Xist* interactors in previous studies^15,16^ and cataloged in the RNA association interaction database (http://www.rna-society.org/raid/). We also identified RNA-binding proteins as potentially novel interactors in this screen (Figure 2e; Table S1). Importantly we demonstrate that specific binding to *Xist* was lost following denaturation of the protein-array (Figure 2e,f; Table S1).

### Target selection, antigen production, and immunization

We initially set out to identify the core set of human proteins that are actively involved in transcriptional regulation. To do this, we integrated previously annotated lists of high-confidence transcription factors (TFs)^17^, along with proteins known or inferred to regulate transcription based on either literature sources or high-confidence direct interaction with annotated TFs (Table S2). This resulted in a final target list of 2229 proteins, of which ~73% were annotated as high-confidence TFs (Figure 3a).

**Figure 3.**
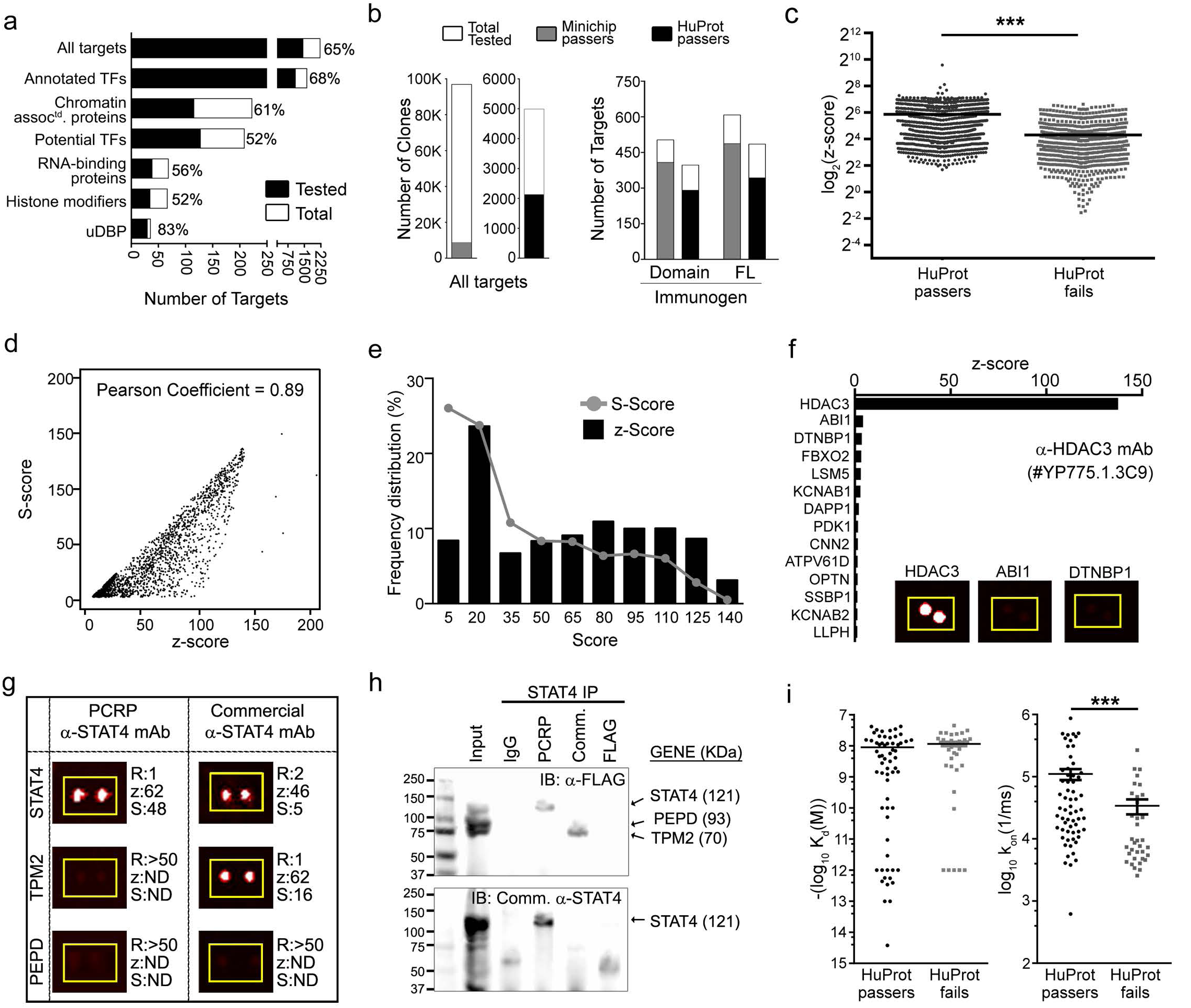
Primary validation of PCRP reagents. (a) The percentage of targets used for immunization is mentioned above bars representing the major target classes. (b) Number of mAbs/targets tested and passed at minichip and HuProt stages. (c) Comparing the HuProt z-score of pass (S-score >2.5) versus fail mAbs. *** represents p<0.0001 by an unpaired t-test with Welch’s correction. (d) For mAbs that pass HuProt, correlation between z-versus S-scores is shown as a scatter plot. (e) Frequency distribution of z- and S-scores amongst mAbs passing the HuProt validation. (f) z-score for targets recognized by the anti-HDAC3 (#YP775.1.3C9) antibody. Insets represents HuProt signal observed for targets indicated. (g) Micrographs representing the HuProt signal observed by either PCRP or commercial anti-STAT4 for the intended target (STAT4) and off-targets (TPM2 and PEPD). Corresponding rank (R), z-score (z) and S-score (S) for each target is mentioned next to the micrographs. (h) IP experiment with either mouse pan-IgG sera (Santa Cruz), PCRP anti-STAT4 mAb (#JH46.1.2A2) or commercial anti-STAT4 mAb (recommended for IP and IB). IP is carried out on a mixture of three lysates from HEK cells transfected with STAT4, PEPD and TPM2. IB is carried out using both rabbit antiFLAG (Cell Signaling) and commercial mouse anti-STAT4 mAbs. (i) Affinity values (K_d_ and k_on_) are compared between mAbs that passed HuProt and those that failed HuProt. *** represents p<0.0001 by an unpaired t-test with Welch’s correction.

This target list was then used to produce recombinant antigens for immunization. A total of 1290 constructs encoding recombinant, His or Halo-tagged domains of individual target proteins were assayed for expression in *E. coli,* and 949 yielded sufficient protein for purification of >0.1 mg of protein (Table S3)^9^. To supplement this antigen supply, 1477 constructs encoding GST-tagged full-length target proteins were expressed and purified from yeast. Of these, 768 yielded >30 μg of purified protein and were used for immunization (Table S4). In summary, we were able to express, purify and immunize 1453 unique proteins (~65%) of our entire target list with significantly varied success among different target classes (Figure 3a).

A total of 646 domains and 668 full-length proteins were used for immunization. For this, we used both systemic intraperitoneal (i.p.) immunization and local footpad (f.p.) immunization, the latter route of administration being used most extensively for proteins that gave low yield. In cases where the immunogen was a GST-tagged full-length protein, an ELISA-based counter-screen was performed to exclude mAbs that recognized GST (~80% of all mAbs tested).

### Analysis of mAb specificity by HuProt binding

Our initial screen for whether IgG+ mAbs could selectively recognize their target consisted of binding assays conducted on a small array comprising 20-80 other antigens used for immunization (minichip). 8.9% of all mAbs tested were found to preferentially recognize their cognate target (Figure 3b). 81.1% of domains and 80.1% of full-length proteins generated at least one mAb that passed minichip screening (Figure 3b). The specificity of a subset of minichip-positive mAbs was then analyzed using HuProt binding. Using criteria established previously^22^ (also see online Methods), a mAb was considered to pass HuProt analysis (HuProt+) if the z-score of the cognate target ranked highest among all 19,030 antigens on the array, with the S-score (or the difference between the first and second-ranked z-scores) >2.5.

Using these stringent criteria, 42.6% of minichip-positive mAbs, corresponding to 73.3% and 70.7% of targets tested using domains or full-length antigens respectively, passed HuProt analysis (Figure 3b). On average, HuProt+ mAbs showed substantially higher binding to their target, as measured by z-score, than did HuProt-negative mAbs, even when only considering cases where the intended target appeared in the top 50 proteins recognized (Figure 3c). We observed a strong correlation between z- and S-scores for passing mAbs, indicating that HuProt+ mAbs showed highly specific binding to their cognate antigen (Figure 3d), with a broad distribution of both scores (Figure 3e). Some mAbs showed exceptionally high S-scores. An example is shown in Figure 3f, with virtually undetectable binding to any proteins other than its cognate target HDAC3.

To determine whether antibody specificity as measured by S-score is a useful criterion for identifying cross-reactivity, we profiled commercially available mAbs that recognized the same target antigen for which we had passing HuProt+ mAb. An example of this analysis is shown in Figure 3g, where mAbs targeting STAT4 are profiled. In this case, while the HuProt+ PCRP mAb was highly selective for STAT4, the commercial mAb recognized TPM2 as its top target and STAT4 as its second target (Figure 3g). The HuProt+ and the HuProt-negative commercial antibodies were then tested by immunoprecipitation, using a cocktail of FLAG-tagged STAT4, TPM2 and PEPD recombinant proteins, a third protein not predicted to be recognized by either mAb (Figure 3h). We observe that the HuProt+ PCRP mAb efficiently and selectively recovered STAT4, whereas the HuProt-negative (HuProt-) commercial antibody recovered TPM2 but not STAT4. This suggests that mAbs displaying high S-scores show high specificity to their intended targets.

Antibodies that are highly specific have been suggested to also bind with high affinity^18, 19^ To investigate whether HuProt+ mAbs also recognized their targets with high affinity, and whether the affinity for their targets was greater than the affinity of HuProt-mAbs for theirs, we measured binding kinetics for 122 HuProt+ mAbs and 106 (minichip-positive, HuProt-) mAbs using a real-time, label-free Octet analysis^20^. Using Kd<50 nM as a cut-off, we observed that 78/122 HuProt+ mAbs and only 44/106 HuProt- mAbs showed high affinity (chi-square test, p<0.001), indicating that HuProt+ mAbs did indeed show higher overall affinity than HuProt-mAbs. Surprisingly, we observed that this difference in affinity was reflected only in the k_on_ scores of HuProt+ mAbs when compared to HuProt-negative mAbs (Figure 3i). However, the z- and S-scores of HuProt mAbs did not correlate with any of the measured affinity parameters [Pearson’s correlation coefficient for z-score and (i) ln(K_d_) = −0.139, (ii) ln(k_on_) = −0.098, (iii) ln(k_off_)= −0.188. Pearson’s correlation coefficient for S-score and (i) ln(K_d_) = −0.214, (ii) ln(k_on_) = −0.072, (iii) ln(k_off_) = −0.23]. This indicates that mAb specificity as measured by HuProt does not serve as a proxy for affinity.

### Secondary validation of mAbs

We considered any HuProt+ mAb, along with a limited number of HuProt-, high-affinity mAbs (Kd<50 nM), as a candidate for secondary validation. In all cases, this involved testing whether the mAbs could successfully recognize their targets using immunoblotting (IB) and immunoprecipitation (IP). We conducted high-throughput analysis of mAbs in these assays (Figure 4a). Full-length target genes were cloned into a doxycycline-inducible expression vector, which were then transiently transfected into Tet-ON HeLa or HEK293 cells^21^. Expression of the target was induced with a low and a high dose of doxycycline to cover a wide range of the target’s expression level in cells. After 24-48 hrs of induction, cells were pelleted for future use or directly lysed for IP analysis. Using either anti-target mAb, anti-FLAG M2 (positive control) or anti-mouse IgG sera (negative control), lysates were immunoprecipitated by an established standard operating protocol (see Methods). The LiCOR two-color infrared fluorescence imaging system was used for simultaneous evaluation of the IP and IB performance of a given anti-target mAb.

**Figure 4.**
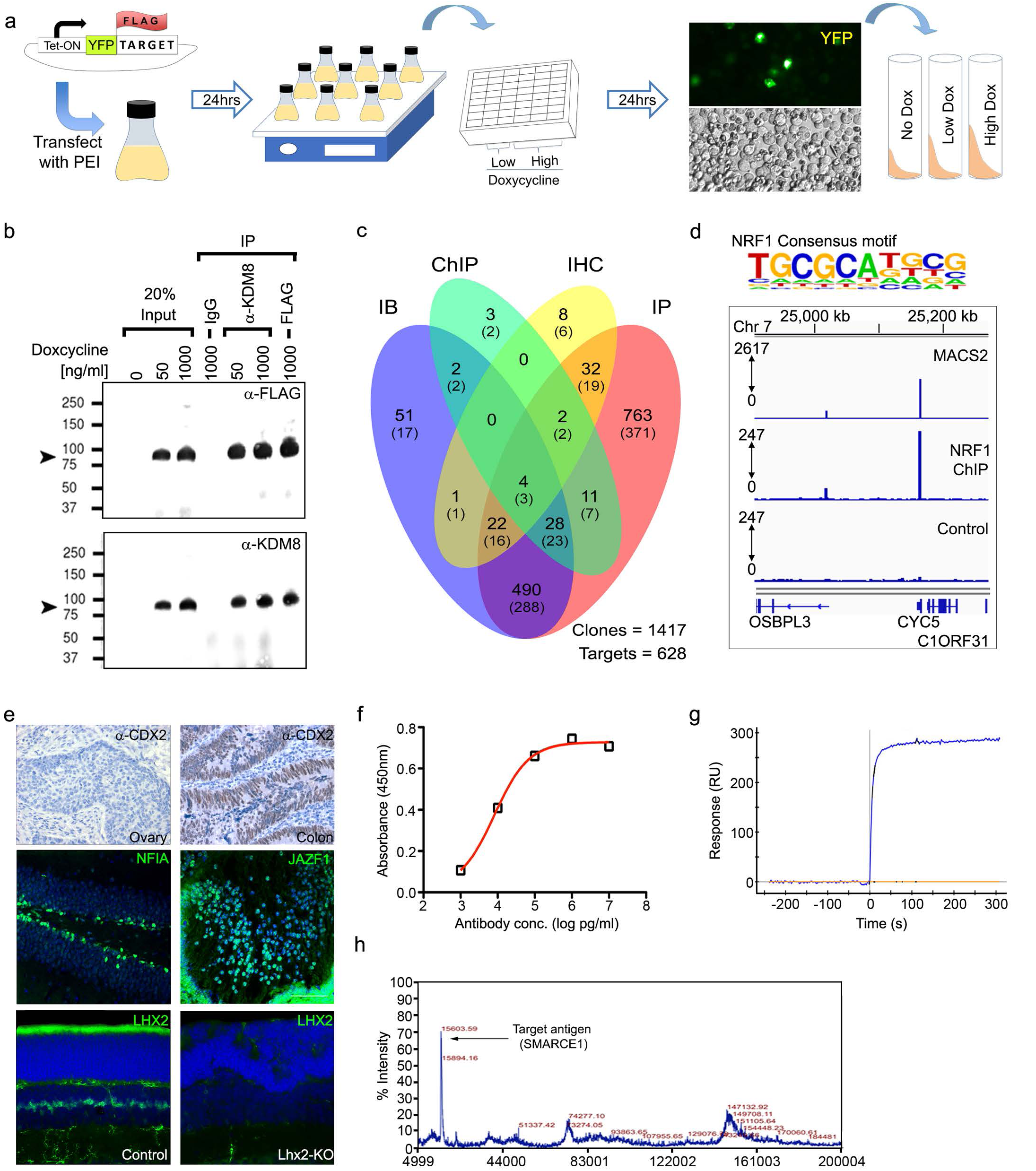
Secondary validation of PCRP reagents. (a) Schematic representation of the high-throughput pipeline for secondary validation. (b) Secondary validation data for a IP+IB+ anti-KDM8 mAb (#YP50.1.1A2). Solid arrow indicates the expected molecular weight of the expressed fusion KDM8 protein (84.6KDa) (c) Venn diagram representing the distribution of mAbs and the corresponding targets in parenthesis, that were empirically determined to be applicable in IP, IB, ChIP and/ or IHC experiments. (d) Representative examples of ChIP-Seq positive mAbs. A known consensus binding site for NRF1 was identified as the top enriched motif in the experiment. A ChIP-seq peak is detected in promoter region of known NRF1 target, CYCS, is graphically represented. (e) Anti-CDX2 (#R1435.1.1A3) exhibits specific nuclear signal in cells of human colon tumors but not in human ovarian tumors. NFIA and JAZF1 expression is visualized in the adult mouse hippocampus (anti-NFIA #R1356.1.2C6) and mouse embryonic (E14) lens (anti-JAZF1 #R913.1.1C2) respectively. P14 mouse pups of the *RaxCreER^T2^;Lhx2^lox/lox^* background were injected with 0.2 mg tamoxifen daily from P1 to P3 to induce conditional deletion of *Lhx2* in Müller glia (Lhx2-KO). Lhx2-KO pups and age-matched controls were processed for IHC with anti-LHX2 (#R911.1.2G5).

For evaluating efficiency of IP, input samples from lysates of +/- doxycycline induction were included (Figure 4b). A given mAb was scored IP+ when anti-FLAG IB showed positive signals at the predicted molecular weight following immunoprecipitation of doxycycline induced samples, but not following immunoprecipitation with mouse IgG. Similarly, a given mAb was scored IB+ when anti-target mAb exhibits positive signal at the predicted molecular weight in any lane except in the uninduced or pan-IgG IP lanes. Evaluation of the performance of each mAb in IP and IB was scored by two independent investigators using a standard set of criteria (see online Methods). Using this validation scheme we identified 763 IP+IB-, 51 IP-IB+ and 490 IP+IB+ mAbs corresponding to 736 targets (Figure 4c).

As our target set of proteins are enriched for transcription factors, we tested the ability of mAbs that are either IP+ and/ or IB+ in chromatin immunoprecipitation (ChIP) application. 305 mAbs to 176 targets were randomly tested first by IB against endogenous targets in selected cell lines. Using ENCODE standards, 50 mAbs against 39/176 targets performed well in ChIP-seq experiments with successful identification of their consensus binding sites (Table S5). For example, 61.22% of targets recovered by ChIP-seq using the anti-NRF1 mAb (#R157.1.3H1) mapped to a known NRF1 consensus site (Figure 4d and Table S5). We performed *de novo* motif enrichment analysis for targets with undefined consensus sequence (Table S5). All of the data generated by ChIP-seq during this project have been deposited to the GEO database.

As transcription factors exhibit exquisite tissue specific expression and are excellent diagnostic markers for tumors, we also randomly evaluated the ability of passing IP+ and/ or IB+ mAbs in immunohistochemistry (IHC) assays. Using a tissue microarray comprised of multiple human tumors, 12/129 mAbs corresponding to 9/73 targets showed positive IHC signals in at least one tumor section (Figure 4e). As expected, the IHC signal intensity correlated perfectly with the profiled mRNA expression levels of the corresponding target in these cancerous tissues^12^. Furthermore, the IHC staining pattern of these mAbs corresponded well with those generated by mAbs considered to be gold standards by pathologists (Figure S1). Next, as TFs share a strong evolutionary conservation between mouse and human, we also independently tested a random set of our passing mAbs on mouse brain and retinal tissue sections using IHC. 58/192 mAbs for 39/93 targets were scored IHC positive by comparing the expression patterns of the targets as described in literature (Figures 4e). As a proof of principle, we further validated the specificity of our mAbs in mouse IHC by utilizing tissues from *Rax-CreER^T2^;Lhx2^lox/lox^* conditional knockout mice^23^. IHC signal using anti-LHX2 mAb (#R911.1.2G5) was completely lost in the *Lhx2* cKO retinal mice, indicating the specificity of our mAbs to the intended target in IHC application (Figure 4e).

To ensure an unbiased evaluation of our mAbs, the PCRP consortium funded an independent body (NIH/NCI) to extensively evaluate a subset of the mAbs that passed our secondary validation pipeline. Three orthogonal approaches were employed: IP-coupled mass-spectrometry (IP-MS), surface plasmon resonance (SPR) and enzyme linked immunosorbent assay (ELISA). Using NCI standard criteria (see online Methods), 43 mAbs to 39 targets were tested and 39, 34 and 22 mAbs passed in ELISA, SPR and IP-MS experiments respectively. Representative examples of a passer in each of these experiments are shown in Figures 4f-h.

### Meta-analysis identifies critical factors that influence generation of passing mAbs

Having generated and characterized a large number of highly specific mAbs against a broad range of proteins involved in transcriptional regulation, we performed a meta-analysis of the data collected during this process (see online Methods for details). First, we observed that the z- and S- scores of mAbs that passed secondary validation were found to be significantly higher than mAbs that failed secondary validation (Figure 5a). However, none of the individual parameters measured significantly correlated with any of the success rates in secondary validation. We therefore explored the alternative hypothesis that a complex interplay of parameters could likely dictate success at different stages of the pipeline.

**Figure 5.**
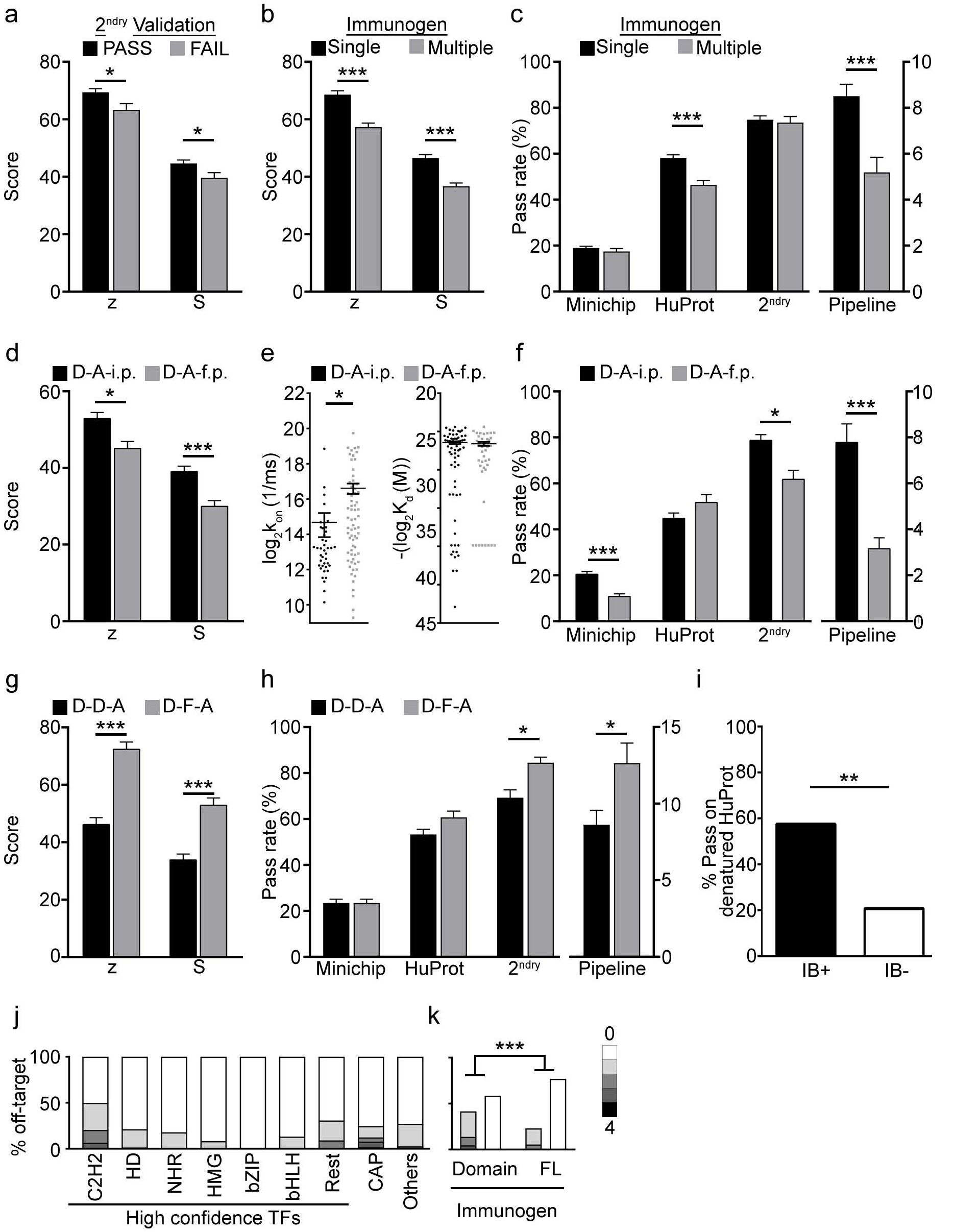
Meta-analysis. (a) mAbs that passed secondary validation have higher z- and S- score. * represents P<0.05 in a unpaired t-test with Welch’s correction. (b-j) *, ** and *** represents p<0.05, p<0.001 and p<0.0001 respectively by Benjamini-Hochberg FDR. (b) Comparing z-/S- score for mAbs raised by immunizing with single and >1 antigen (multiple). (c) Comparing success rates at different stages of the pipeline for mAbs raised by immunizing with single and >1 antigen (multiple).(d-h) key to the graphs represent abbreviations for mAb grouping (see online methods for details). D-A-f.p. = Domain-All-footpad. D-D-A = Domain-Domain-All. D-F-A = Domain- full length-All. (d) z-/S-scores of mAbs generated by i.p is statistically higher than f.p. (e) k_on_ but not K_d_ is significantly lower in mAbs raised by i.p. Vs f.p. (f) Success rates at different stages of the pipelines for mAbs. Overall pipeline performance of mAbs immunized by f.p. is significantly lower compared to i.p. (g,h) Within mAbs generated using a single domain immunogen, mAbs that recognize their cognate full-length (FL) target have higher average z-/S-scores, and better success rates throughout the pipeline when compared to mAbs that only recognize their cognate domain immunogen. (i) 27/47 IB+ mAbs pass screening on denatured HuProt while only 6/29 IB-mAbs pass this screen. **represents p<0.001 in Fisher’s exact test. (j) Identifying the percentage of off-targets (rank 2-5 with z-score>3) for each individual mAb that belong to the intended target’s class/subclass. The fraction of mAbs in which all off-targets do not belong to the same class/subclass as the intended target is represented by the unshaded white portion. Light grey to black shades represent fractions wherein one through all four off targets belong to the same class/subclass as the intended target. (k) This distribution in off target’s class/subclass is also represented based on the type of immunogen used for generating the mAb. *** represents p<0.001 in a Fisher’s exact test.

Detailed immunization records for a subset of targets were not adequately maintained and were therefore excluded from all further analysis. We performed three types of parametric comparisons at the level of either: (a) individual mAbs (Table S6), (b) targets (Table S7) or (c) target sub-classes. Matrix comparison of parameters (Table S8) generated a comprehensive list of differentiating factors that potentially identify success in the pipeline (see Tables S9-14). We manually selected comparisons to exclude potential bias due to skewed sample size, or in cases where interpretation was difficult (see online Methods).

We had hypothesized that immunization with pools of disparate antigens might increase the probability of recovering high quality mAbs, at least to a subset of the targets. However, parametric comparison clearly shows that mAbs generated by immunization with a single antigen exhibits better average z- and S- scores (Figure 5b and Tables S9: comparisons #2 and #17; p=6.52E-09 and 6.78E-09 respectively) and significantly improved success at the HuProt stage of the pipeline (Figure 5c and Tables S10: comparisons #12; p=4.18E-07).

We grouped mAbs based on procedures used to generate the mAbs (see online Methods for detailed explanation). These groups are represented by abbreviations separated by hyphens in the following order: (i) type of immunogen used - domain (D), full-length (FL) or all (A) - (ii) type of epitope recognized by the mAb - its immunized domain but not full-length counterpart on the HuProt (D), both its domain immunogen and full-length counterpart on HuProt (FL) or considering all types (A) - (iii) route of immunization-intraperitoneal (i.p.), footpad (f.p.) or all immunizations (A). Therefore, the D-A-i.p group refers to mAbs generated using a domain immunogen, that recognized either its cognate domain immunogens in our screens and/ or their cognate full-length target on HuProt, generated by i.p. immunization route. Similarly, D-D-A group refers to mAbs generated using a domain immunogen, that recognized their cognate domain immunogens in our screens, but not their cognate full-length target on HuProt, generated by any of the immunization routes used during the course of this project. For more detailed description kindly refer to online Methods section.

We compared mAbs in the D-A-i.p and D-A-f.p groups generated by using both single and multiple immunogens. Immunization by i.p. consistently generated mAbs that exhibited statistically higher z-, S-scores when compared to f.p. Immunizations (Figure 5d and Table S11:comparisons #2 and #10; p=3.06E-03 and 6.00E-05 respectively). Although no differences in K_d_ were observed, mAbs generated by f.p exhibited a statistically higher k_on_ score than those generated by i.p. (Figure 5e and Table S11:comparisons #34 and #18; p=9.82E-01 and 6.7pE-03 respectively). Furthermore, i.p. immunization gave a substantially higher fraction of passing mAbs, at minichip and secondary validation stage, than did f.p. immunization (Figure 5f and Table S12:comparisons #2 and #18; p=1.17E-06 and 7.11E-03 respectively), indicating the i.p. route to be the best choice for immunization. However, this observation needs to be reconciled with the fact that the concentration of proteins injected by i.p. route is, on average, two orders of magnitude higher than for f.p., and the loss in efficiency is easily offset by the economic savings in antigen production.

Next, we compared parameters and success rates measured for the D-D-A and D-F-A groups. To maintain sample homogeneity, we focused comparison on only mAbs that were generated using a single immunogen. Strikingly, D-D-A mAbs exhibited significantly lower z- and S-scores on average (Figure 5g and Table S13: comparisons #5 and #27; p=1.83E-13 and 1.08E-08 respectively). These differences were also visible at the level of targets. D-F-A mAbs had higher success rates in secondary validation and therefore in the overall pipeline (Figure 5h and Table S14: comparisons #43 and #62; p=7.11E-03 and 5.01E-03 respectively). Taken together, these observations suggest that mAbs that recognize their intended full-length targets on HuProt are superior in almost all measured parameters.

### Analysis of denatured HuProt identifies IB-grade mAbs

As demonstrated in Figure 2, proteins on the HuProt are primarily in native conformation. Furthermore, only 3.6% of passing mAbs were IP-negative and IB-positive. This implied that mAbs identified in our HuProt analysis preferentially recognize native epitopes, and that denaturing the proteins on the HuProt could selectively identify IB-grade mAbs. To test this hypothesis, we determined z- and S-scores for both IB+/IP+ (47 mAbs to 38 targets) and IB-/IP+ (29 mAbs to 28 targets) using denatured HuProt arrays (Table S15). As expected, we observed that IB+ mAbs passed our denatured HuProt screen (rank1 for intended target and s-score >2.5) more often than IB-negative mAbs (Fisher’s exact test, p=0.002) (Figure 5i).

### mAb specificity is influenced by the source of immunogen

For mAbs that had passed our validation criteria (S>2.5 and IP+ and/or IB+), there are a subset of mAbs that bound to at least 4 potential off-targets (rank 2-5, z-score >3) albeit with lower specificity than the intended target. We analyzed this subset of mAbs and found that on average ~71% of all off-targets are not in the same target class/subclass as the intended target (Figure 5j). Interestingly, off-targets of mAbs generated using domains are more likely to be in the same target class/subclass as compared to those generated by full-length (FL) antigens (Figure 5k; Fisher’s exact test, p=2.84E-35). Our results suggest that screens for mAb specificity that analyze only close homologs of the intended target will not identify the majority of cross-reactive proteins.

## Discussion

This effort represents the largest systematic attempt to date to generate and extensively validate a collection of renewable affinity reagents directed against human proteins. A summary of the reagents generated through this production and analysis pipeline is shown in Figure 6. As discussed in Figure 5, we list all the mAbs (Table S6), their corresponding targets (Table S7) by their class/sub-class (Tables S16 and S17) and success rates across the pipeline by the type of immunogen and route of immunization used (Table S18). A total of 1406 IP and/or IB-grade mAbs were obtained against a total of 736 target proteins.

**Figure 6.**
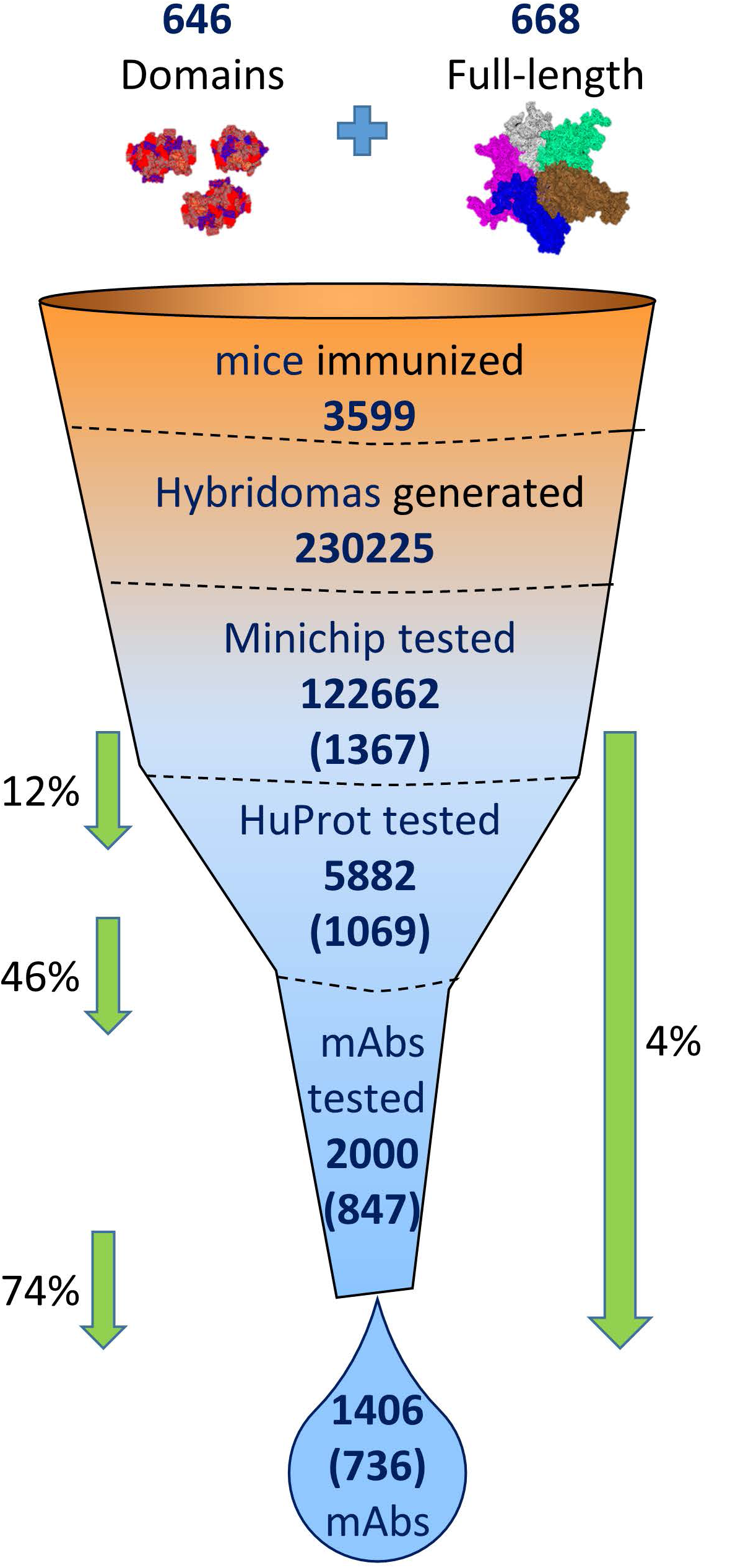
Summary analysis of the PCRP pipeline. A schematic representation highlighting the attrition funnel through which immunogens entering the pipeline ultimately generate high-quality mAbs. Number of mAbs and targets (mentioned in parenthesis) that were tested are indicated for each step of the validation pipeline. Overall pass rate for the entire pipeline at minichip, HuProt, secondary validation and in the overall validation are indicated.

The vast majority of our mAbs (96.4%) were IP-grade, with many also usable for IB. A variety of functional tests clearly showed that proteins on the HuProt array are predominantly in native conformation (Figure 2), and that mAbs identified as specific on HuProt are therefore likely to perform robustly in IP analysis. We also showed that mAbs that recognize linear epitopes, as determined by success in IB analysis, were more likely to selectively recognize their cognate target on denatured HuProt arrays (Figure 5i). This suggests that array-based screening can be adapted to identify antibodies based on their desired application. However, a substantially smaller fraction of our mAbs is suitable for IHC or ChIP-Seq (Figure 4c). In fact, stringent post-hoc evaluation of IP/IB performance of mAbs led to the labelling of some mAbs as non-passers (IP^-^IB^-^); some of these same IP- IB- mAbs however, had been independently assessed as passers in ChIP-seq (3 mAbs to 2 targets) and IHC (8 mAbs to 8 targets). These observations indicate that mAbs that do not pass HuProt analysis may nonetheless be usable in certain contexts, particularly if cross-reacting proteins are neither expressed nor accessible in the cell or tissue type under investigation. In line with our demonstration that screening mAbs against a denatured HuProt (Figure 5i) leads to the preferential identification of IB+ mAbs, it is likely that mAbs counter-screened against HuProt pretreated with detergents, fixatives or solvents used in specific downstream applications such as immunocytochemistry, might enrich for mAbs that perform well in those applications.

The collection of mAbs generated and described here is currently distributed through both DSHB and commercial suppliers. Our mAbs have a number of unique advantages compared to resources generated by similar efforts. First, these reagents are renewable, in contrast to collections of polyclonal antibodies. Second, since they were generated using recombinant domains and full-length proteins and screened using native HuProt arrays, they are highly enriched for IP-grade mAbs. This stands in contrast to other collections generated using linear peptides and protein fragments, which generally work well in IB and immunostaining, but fail to reliably recognize target proteins in native conformation. Third, the use of animal-based methods, as opposed to purely *in vitro* screening, increases the likelihood of identifying research-grade reagents by enriching for reagents that recognize a mixture of linear and conformation-specific epitopes.

The costs of generating and validating research-grade antibodies are high, with estimates averaging ~$40,000/research-grade reagent, and can be higher for recombinant antibodies. Most of these costs are incurred during the process of validation^6^. Using the production pipeline described here, we achieved a substantial reduction in cost, with average cost/mAb at $7,500 per passing reagent. A major bottleneck for all antibody production efforts is antigen production, particularly when large quantities of native proteins are required^24^. This also proved to be a major problem in early phases of this project. To a large extent, we were able to overcome this problem by implementing footpad immunization, which requires substantially less antigen than intraperitoneal injection. While full-length protein production in yeast gives substantially lower yields than protein production in *E. coli,* immunization of full-length antigens by the footpad route yielded mAbs that are superior in z-/S- scores (Figures S2a and Table S13: comparisons #1 and #23; p=3.65E-06 and 1.01E-02 respectively) and exhibited comparable success rates as mAbs generated by immunization of domains by this route (Figure S2b and Table S14; comparisons #2,21,40 and 59, p>0.05).

Our meta-analysis reveals that HuProt screening is highly predictive of success in secondary validation and screens out mAbs with cross-reactivity to all classes of proteins. Indeed, analysis of commercially available mAbs on HuProt was found to predict cross-reactivity against previously unsuspected targets (Figure 3g,h). Furthermore, array-based analysis can be a highly cost-effective approach for both production and identification of highly specific antibodies. The low cost and high-throughput features of this production pipeline renders it potentially suitable for generating renewable affinity reagents to many other protein families, and implies that a similar effort aimed at targeting the remainder of the annotated human proteome is feasible and economically viable. Lastly and most importantly, there is finally a growing awareness of the antibody reproducibility crisis^1-7^, and concerted efforts to define a global standard for antibody validation criteria are being made^25-27^. Our study demonstrates the clear need for using a comprehensive protein array, like the HuProt, as a first pass to prove mAb specificity, and demonstrates the broad usefulness of protein array-based specificity analysis as a standard approach for antibody validation.

## Online Methods

### Validating and Curating clones used in HuProt

Proteins spotted on HuProt were purified from yeast transformed with expression vectors encoding the said ORF^1^. To generate detailed annotation of the ORFs we Blast+ aligned the ORF sequence to multiple public databases (UniProt, CCDS, RefSeq, Ensembl) to generate an integrated alignment score for each of our clones. If a clone covered the entire sequence of a known protein we would consider the clone as a full-length (FL), while partial matches were regarded to truncated (TRUNC) clones. As our source clones were derived largely from ORFeome collection repositories, it included ORFs containing UTRs, unannotated splice-variants and SNPs. To account for this diversity, we developed numerous quality control (QC) parameters to categorize our clones ranging from perfect matches to the known proteincoding transcriptome to yet unreviewed potential protein-coding ORFs. Detailed breakdown of this classification along with our threshold parameters can be accessed at https://collection.cdi-lab.com/public.

Having annotated our ORF collection, we next sought to comprehensively validate the integrity of the collection. Our yeast expression vectors were maintained in *E.coli* bacterial stocks. Historically QC was performed by traditional restriction digestion of the construct and gel-electrophoresis to confirm the presence of the correct sized ORF. To systematically check and confirm identity of each of the ORFs, we performed bi-directional Sanger sequencing (Beckman-Coulter Genomics) to cover ~800-1000bp each at the 5’ and 3’ ends of the insert. Sequences were first trimmed to represent only high-quality reads (phred >20) represented on average by the central 700bp of a given read. We used Blast+ to align reads to the expected record ORF sequence and also to the CCDS database to find the best matches. We would consider a pair of reads as ‘PASS’ if at least one of the reads matched the reference in our records, with less than 3 substitutions, and no deletions or insertions. Surprisingly about 8% of our collection was found to contain inconsistencies as per sequencing and were therefore retired from any further HuProt analysis. These failed clones included instances wherein forward and reverse reads either matched to different references (labelled as ‘Mixture’) or matched perfectly to an ORF that was unrelated to the one in our manual records (labelled as ‘Mislabeled’). Some clones had lost their inserts and instead aligned to the vector sequence (labelled as ‘Empty’) or simply did not align to any known sequence (labelled as ‘No Match’ or ‘Sequencing FAIL’).

### Target selection

In addition to the high confidence transcription factors (TF) annotated previously^1^, the list of proteins with TF-like activity was inferred using relevant GO annotations such as “regulation of transcription, DNA-dependent”. Other genes that were well-established transcriptional coregulators or epigenetic modifiers based on the literature, but not classified as such by GO, were added manually. Unconventional sequence-specific DNA binding proteins (uDBPs) that showed clear evidence of being able to bind specific DNA sequences by EMSA^2^ were also added. Finally, TF-associated proteins were identified from protein-protein interaction (PPI) databases and a subset of these were included, prioritizing only those considered to be both of high interest to the scientific community and lacking high-quality commercially-available reagents.

### Antigen production

TF antigens used for immunization, in the forms of domains or full length proteins, were produced in two expression systems. Defined TF domains were expressed as AviTag-labeled proteins from a T7-containing plasmid in *E. coli* (Tuner(DE3)/pLysSRARE2) grown in LB with appropriate antibiotics. Protein expression was induced with IPTG, cultures were grown for 16 hr at 17°C, and pelleted cells were lysed by sonication in 50 mM Tris-HCl, 6 M urea, 10 mM imidazole, 500 mM NaCl, 1 mM TCEP, 0.02% NaN3, pH 7.5. Lysates were clarified by centrifugation (30 min at 10,000 × *g*, 4°C), and the soluble fraction was used for HisTrap-mediated purification. Eluted proteins were dialyzed in 20 mM HEPES, 100 mM NaCl, 400 mM L-arginine, 20% glycerol, 1mM TCEP, 0.02% NaN_3_, pH 7.5. Purity was assessed by SDS-PAGE and MALDI-TOF MS, and if necessary proteins were further purified by gel filtration chromatography in the same buffer. At early stages of the program, antigens were suspended in PBS to reach the desired concentration for immunization. However, when it became clear that high concentrations of arginine and/or NaN_3_ were causing footpad necrosis, antigens were then concentrated, dialyzed 3x against PBS containing 10-20% glycerol, and in some cases supplied as a mixed suspension of soluble and precipitated material. Antigens were then stored frozen at -80°C until needed for immunization.

Full length transcription factors were expressed as GST-fused proteins from a galactose-inducible, Ura+ plasmid^1^ in S. *cerevisiae* that was grown in SC/ura-/raffinose medium. Protein expression was induced with galactose, cultures were grown at 30°C for 18 hr, and pelleted cells were lysed by vortexing at 4°C with zirconium beads in 50mM Tris, 100mM NaCl, 1mM EGTA, 10% glycerol, 0.1% Triton X-100, 1X Complete (Roche) protease inhibitor cocktail, 0.1% b-mercaptoethanol, 1mM PMSF, pH 7.5. Expressed proteins were purified with glutathione-sepharose, and the eluted proteins were concentrated to at least 1 mg/mL with spin-concentrators and stored frozen at -80°C until needed for immunization.

### Animal work and immunization

For immunization of mice, antigens were adjusted to a concentration of 2-4 mg/mL by spin concentration or by dilution with PBS. For footpad (f.p.) immunization, 25-30 μg of protein in 12.5 μL were mixed with an equal volume of TiterMax adjuvant and vortexed. The complete emulsion mixture for each protein was injected into one rear footpad of a 6-8 week old female BALB/c or CD-1 mouse. In some cases, mice were euthanized 12-14 days later and popliteal lymph nodes aseptically collected. In other cases, twelve days after footpad immunization, an equal amount of antigen/adjuvant was injected into the hock area above the same foot. Three days later these mice were euthanized and the popliteal lymph nodes were aseptically harvested for hybridoma production. For intraperitoneal (i.p.) immunizations, 100 μg of antigen was mixed with an equal volume of Sigma Adjuvant system adjuvant and vortexed. The entire mixture was injected i.p. on day 0. On days 14, 28 and 42 the same amounts of antigen/adjuvant were administered i.p., mice were euthanized on day 46 and spleens were aseptically harvested for hybridoma production. For a small cohort, a combination of i.p. and intravenous (i.v.) injection was used. On day 0, 100 μg antigen plus adjuvant (Sigma Adjuvant system) were injected IP, and on days 14 and 21, 50 μg antigen mixed with adjuvant were injected IP. On days 28, 29 and 30, 50 μg of antigen without adjuvant were injected into the tail vein. On day 31 the mice were euthanized and their spleens aseptically harvested for hybridoma production.

### Hybridoma generation

Immune lymph nodes and spleens were teased into single cell suspensions in DMEM containing 100 U/mL penicillin and 0.1 mg/mL streptomycin (P-S), and cell numbers and viability were determined. Immune cells and Sp2/Ag0 myeloma cells (ATCC) were washed 2 times with DMEM/P-S, combined at a 5:1 ratio, and fused with 50% buffered PEG (Hybri-Max, Sigma), following standard procedures^2^. Fused cells were plated in DMEM/20% FBS/P-S/1X HAT/1X HFCS/2% methylcellulose/1 μg/mL goat anti-mouse IgG-AlexaFluor 488. Following growth for 8-10 days at 37°C, 5% CO_2_, plates were scanned with a fluorescence dissecting microscope (Olympus MVX10), and positively staining colonies were transferred with microcapillary pipets into individual wells of 96-well plates. Colonies were expanded in DMEM/10% FBS/P-S/1X HT, at which point supernatant was assessed for the presence of IgG by ELISA. Supernatants from IgG-secreting clones were tested by GST-specific ELISA to identify and eliminate clones producing antibodies that bound to the fusion tag.

### Primary validations

To identify antibodies that bound to the target TF immunogen, microarrays that were created on glass slides by spotting the immunogens in a 2 × 7 sub-array pattern (“minichips”) on epoxide-coated slides using an ArrayIt NanoPrint™ LM210 microarray printer. Slides were fitted with 2 × 7 format gaskets, blocked with PBS/BSA, and IgG-positive culture supernatants were added to individual sub-array wells. Binding of an antibody to the spotted proteins was revealed by incubating all wells with goat anti-mouse IgG (H+L)-AlexaFluor647, washing, and scanning the arrays with a Molecular Devices GenePix 4000B scanner supported with GenePix Pro 7 software. A positive signal was considered one to be significantly brighter than all other spots in the same well when examined by eye. Clones secreting antigen-binding antibodies were expanded to T-600 flasks, and the cells were collected by centrifugation, suspended in growth medium plus 10% DMSO, and frozen in liquid nitrogen. The associated culture supernatants were used for purification of IgG with protein G-sepharose. To determine the relative level of target protein binding specificity, the purified IgGs were individually used to screen HuProt arrays that contained >17,500 pairs of purified human proteins. Antibody binding to its cognate target on the HuProt was scored using the z- and S- scores as described previously^3^. Briefly, the z-score represents the strength of a signal that a mAb (in combination with a fluorescently-tagged anti-IgG secondary antibody) produces when binding to a particular protein on the HuProt array. z-scores are described in units of standard deviations (SDs) above the mean value of all signals generated on that array. If targets on HuProt are arranged in descending order of the z-score, the S-score is the difference (also in units of SDs) between the Z-scores of consecutive targets. S-score therefore represents the relative target specificity of a mAb to its intended target. A mAb is considered to specific to its intended target if the mAb has an S-score of at least 2.5.

### Cell culture for secondary validation

Adherent 3G-Tet-ON HeLa cells (Clontech #631183) was maintained in DMEM (high glucose, GIBCO) supplemented with 10% FBS (tetracycline free, Clontech #631106), 4 mM Glutamine (Gibco) and 100 units/ml penicillin/streptomycin (Invitrogen, Carlsbad, CA).Tet-On HEK293_LD_ suspension cell^5, 6^ were grown in Freestyle 293 medium (GIBCO) supplemented with 4 mM L-glutamine and 1% FBS (tetracycline free, Clontech #631106) in 8% CO2. Suspension cells were maintained between 0.3-3 × 10^6^/ml in disposable baffled and vented PETG shake flasks (Thermo-Fisher) at 165 rpm using Celltron orbital shaker (25 mm orbital diameter, INforsHT). Suspension cells were always seeded at a volume up to 30%–40% of the culture bottle’s capacity.

### Plasmids and transfection

We used a Gateway-compatible and tetracycline inducible human expression vector (HuEV-A) that was described before^6^ for expression of the intended target of a test antibody. The HuEV-A (Addgene Plasmid #68342) construct appends a N-terminal fusion of 3x Flag peptide, a V5 epitope tag followed by Venus fluorescent protein (YFP variant) to the expressed protein. Transfection of expression constructs in the adherent HeLa cells was performed using FugeneHD (Promega) as per manufacturer’s instructions. For suspension cells, transfection was carried out using Polyethylenimine “Max” high potency linear PEI (Polysciences) as described before^7^. We observed a target dependent sensitivity of certain proteins to form aggregates when overexpressed following induction. By titrating down the doxycycline levels to reduce induction of the protein, and by reducing the incubation time post-transfection, the majority of such aggregation artifacts were overcome.

### Immunoprecipitation and Immunoblotting

16-24 hrs post-transfection, cells were split into six fractions - one fraction was left uninduced as control, two fractions were induced with low levels of doxycycline (1-50 ng/ml) and the remaining three fractions were induced with high levels of doxycycline (20-1000 ng/μl). Each fraction corresponded to ~5×10^5^ adherent HeLa cells or ~2×10^6^ HEK293_LD_ suspension cells. After 16-48 hrs of induction, expression of target protein was confirmed by screening for YFP expression. Following confirmation of YFP expression, cells were harvested in PBS, pelleted, and stored at - 80°C till further use. On the day of immunoprecipitation (IP), respective pellet fractions of a given target were vortexed in lysis buffer (100mM Tris-HCl, 150mM NaCl, 25mM NaF, 50 μM ZnCl_2_, 15% glycerol, 1% Triton X-100) supplemented with protease inhibitors (Roche #11697498001) and BitNuclease (Biotools #B16003). Following a 30 min incubation at 4°C, ~80% of the supernatant was collected after centrifuging at 4000 rpm for 20 min at 4°C. An aliquot of the supernatants were stored as input for the IP.

Supernatants from samples induced with high levels of doxycycline were next incubated with 5 μg each of either mouse IgG sera (#sc-2025, Santa Cruz), the test CDI monoclonal antibody or 1 μg of the positive control monoclonal M2 anti-FLAG antibody (#F1804, SIGMA). Additionally, supernatants from samples induced with low levels of doxycycline were also incubated with 5 μg of the test CDI monoclonal antibody, to test the sensitivity range of the antibody in IP experiments. Incubation was carried out with gentle rocking, overnight at 4°C. Next, the antibody-protein complexes were pulled down by incubating 2 hrs with Protein G dynabeads (Life Technologies #10004D) at 4°C, washed three times with lysis buffer and eluted in 35 μl LDS-sample loading buffer supplemented with *β*-mercaptoethanol as reducing agent (Life Technologies #NP0008). Input lysate, along with the IP samples, were resolved in a SDS-PAGE gel and immunoblotted using both rabbit anti-FLAG (1:1000, #2368, Cell Signalling) and the test monoclonal antibodies from CDI. Anti-rabbit IRDye680RD (LiCor # 925-68071), Anti-mouse IRDye800CW (LiCor # 926-32210) and light-chain specific anti-mouse AlexaFluor 790 (Jackson Immunolabs #115-655-174) secondary antibodies were used for visualization of bands, and the blots were imaged using an infra-red fluorescence imager (LiCor Clx).

### Immunohistochemistry (IHC)

IHC on human tissue was performed on five micron thick sections, cut from formalin-fixed, paraffin-embedded human tumor specimens and mounted on charged slides (Mercedes Medical). The slides were deparaffinized in SlideBrite, washed with 100% ethanol, followed by rehydration in a graded ethanol series (95%, 70% and 50%). Endogenous peroxidase was blocked by immersing slides in 3% hydrogen peroxide for 5 min followed by washing in distilled water. Antigen retrieval was performed in 10mM citrate buffer (pH 6.0) by heating slides in a pressure cooker (Decloaking Chamber, Biocare Medical) at 125°C for 20 minutes. Tissue slides were cooled to 80°C and washed in water. All primary antibodies were diluted to 2ug/ml in 10mM PBS, pH 7.4, containing 3% BSA and 0.05% azide. Slides were incubated for 30 minutes with the diluted antibodies and then rinsed with PBS containing 0.05% Tween-20. A goat anti-mouse horseradish peroxidase polymer detection (ScyTek Laboratories) was applied to tissue sections for 15 minutes at room temperature and rinsed in PBS as before. Sections were finally incubated in 3,3’-diaminobenzidine (Scytek Laboratories) for 5 min. Counterstaining was done with hematoxylin. Negative controls consisting of diluent with no antibody were used in all experiments.

IHC on mouse retina and brain tissue was carried out as described previously^10,11^. Briefly, tissue was collected after perfusing animal, post-fixed after harvest in 4% paraformaldehyde for 1 hr at 4°C or fresh frozen in Clear Frozen Section Compound (VWR), and stored at- 80°C till used for cryo-sectioning. Glass slide mounted tissue section (15-20um sections) are first blocked with standard blocking buffer and then incubated with primary antibodies (1:200) overnight @ 4°C. After washing with 1× PBST (1× PBS+ 0.05% Triton X-100), secondary antibodies conjugated to AlexaFluor 488 (1:500) were incubated for 2 hrs at room temperature in the dark. Subsequent to washing and counterstaining with nuclear stain, DAPI (1:5000), slides are mounted using VectaShield (Vector Laboratories) and recorded using confocal and/or BZ-X700 Keyence fluorescence microscope.

### Image processing and quantification

The software was developed to allow faster and more automated processing and analysis of gel images. It consists of a web-based interface and server-hosted Python and Perl scripts for data organization and image processing. The left pane of the interface provides a tree-view of all uploaded files and allows for creation of folders for organization. The right pane is the analysis window. The initial input for the software is a High Dynamic Range Image (HDRI) in red/green channels, for which the software will convert to a format that can be displayed on screen, and a text file with input parameters. The image is of a ‘gel collage’ – a set of gels arranged on a single image. The software contains a custom algorithm to detect and separate each gel in the collage image. The interface provides a proposed sectioning scheme, which can be adjusted by the user and then saved. Once saved, the gels are separated into individual images and each image can be selected for further processing.

For each gel, the software will automatically detect bands in the marker lane and associated masses, and their location will be indicated on screen for the user to adjust manually, as needed. These locations will be saved and used later in the processing pipeline to create a labeled image. The software also allows for adjustment of the clarity of the image by allowing the user to interactively set the minimum and maximum display pixels for the conversion from HDRI. Further, the software provides an automatic densitometry calculation. The lanes and relevant bands are detected by a custom algorithm and band intensity is measured from the original HDRI image pixels. The signal is calculated as sum of intensity minus background, where background is the median intensity over a 3-pixel wide rectangle surrounding the band. The user may adjust the outline defining the detected bands interactively. All densitometry calculations are then saved to an output text file. The software can also produce labeled images for catalog use, which is currently customized for CDI Laboratories and based on the adjustments made by the user and the input parameter file. These labeled images are made available to download as a zip file via a link in the web interface.

### Meta-analysis of pipeline efficiency

We empirically determined z- and S- scores for mAbs that were tested on HuProt, and for a select subset also measured affinity values by Octet and/or OIRD. We compared differences in procedures used to generate the mAbs and/ or structural differences among the intended targets of individual mAbs. Based on the procedures used, our mAbs could be categorized into groups based on four variables - (a) the type of antigen used (domains (D) vs full-length (FL) protein), (b) whether or not a mAb raised against domains recognized only the immunized domain (D), or both the domain and its full-length (FL) counterpart on the HuProt, (c) the immunization protocol (i.p. vs f.p.) used and (d) whether a single antigen or a pool of antigens was used for immunization. Unless mentioned otherwise, we made comparisons to mAbs generated using both single and multiple immunogens. Furthermore, as detailed immunization records for a subset of targets were not adequately maintained during the initial phase of the project, these were excluded from further analysis, unless mentioned otherwise. By combining the remaining three variables, we categorized our mAbs and represented them using the single letter abbreviations mentioned above. The letter “A” was used when all sub-groups in a given variable were considered. A sample list of such combination is presented below:

| Abbreviation used | Combination represents mAbs generated using- |
| --- | --- |
| D-D-A | Domain immunogen, that recognizes its cognate domain immunogen on HuProt, generated by any of the immunization routes. |
| A-F-f.p | All immunogen types, that recognizes the intended full-length target on HuProt, and was raised using footpad immunization |
| D-A-i.p | Domain immunogen, that recognizes either its cognate domain and/or full-length target on HuProt, generated by any of the immunization routes. |

At the structural level, targets are hierarchically classified wherein multiple targets are represented in each class/subclass, and multiple mAbs represent each target.

Success rate calculated at the level of target, target-subclass or target-class is defined as the fraction of corresponding mAbs that passed relative to the total number tested. Success rates were calculated at the stages of minichip, HuProt, secondary validation, and in the overall pipeline (by multiplying success rates of minichip x HuProt x secondary validation).

We wished to generate comparisons by combining the procedural groups with structural classification. If a target was represented by mAbs generated by more than one type of procedure, we separated them into a category labelled “mixed”. We restricted comparisons of groups with at least 5 mAbs and excluded a subset of comparisons manually to remain conservative in making conclusions. Two prominent examples of such exclusion are (1) mAbs generated from FL antigen using i.p. injection, as this represented a very small subset and (2) comparisons with mAbs in the ‘mixed’ category. P-values were calculated using the statistical software R, by using the t-test function, and effect size was calculated using cohensD function of package ‘lsr’. Benjamini-Hochberg FDR was calculated using R p.adjust function. For FDR calculations, we grouped hypotheses into two groups representing comparisons made on parameters measured for individual mAbs (Tables S9, S11 and S13) and at the level of targets (Tables S10, S12 and S14).

### Scoring and threshold criteria for mAbs

For secondary validation, antibodies were evaluated using preset and standardized thresholds for quality and efficiency in immunoprecipitation (IP) and immunoblotting (IB) applications. Listed below are the parameters that were considered when establishing the thresholds of these validations. To evaluate an IP-grade mAb, following densitometric analysis of signal intensity in the immunoblots, test antibodies with greater than 0.15% IP-recovery, as compared to the total input sample, were considered as passers. Results for each antibody were manually inspected by at least two independent investigators before being passed. We observed target-dependent variance in target stability during the transfection, storage and/or during the IP experimental procedure. As long as the target protein at the expected molecular weight was observed in the input and corresponding IP lanes, the reagent was considered to pass. The ability of the antibody to IP-recover the degraded fragments was not evaluated or considered towards the efficiency of the IP-recovery.

All constructs expressing the target ORF were verified for integrity by Sanger sequencing the construct expression cassette. However, we observed up to 20 kDa differences relative to the expected molecular weight for certain target proteins in our SDS-PAGE gels. As the protein was not visible in the lanes containing lysates of uninduced cells, we considered such cases to be under the normal purview of aberrant migration reported for many proteins in the literature.

As we tested our IP experiments in Tet-ON HeLa or HEK cell lines, we expected to observe no expression of the transfected protein in the absence of tetracycline/doxycycline treatment. However, we occasionally observed leaky expression of the target protein even in the absence of doxycycline, a known property of the older generation of Tet-ON expression cassettes used in our cell-lines. We considered the experiment to be acceptable if the levels of target protein visibly increased at least 2-fold when comparing the uninduced and highest induction samples in the input lanes.

For unknown reasons, in a small subset of our experiments we observed that the rabbit anti-FLAG mAb (Cell Signaling) recognized non-specific signal(s) in all three input lanes. This band intensity did not change as a function of doxycycline treatment, and was not recovered by IP using the IgG or test antibodies. Such non-specific signal was not considered to be relevant for experimental evaluation of the antibody. We observed that certain target proteins had an increased propensity to form insoluble/”sticky” protein aggregates when expressed at high levels. This feature was readily visible as a strong signal in the IgG-IP lanes of our experiments. By serially titrating the level of induction by doxycycline, we were able to prevent this unwanted aggregation of target proteins with concomitant loss in signal intensity of the IgG-IP lanes. However, in certain scenarios a weak but visible signal might be present in in the IgG lane. If the densitometric signal observed in the IP lane (lane 6) using the test mAb was found to be at least two-fold higher than the cognate signal in the IgG-IP lane (lane 4), we considered such an antibody to be a passing reagent.

In a significant fraction of our experiments we observed failure of recovery in our positive control using the mouse anti-FLAG IP (lane 7). As the FLAG tag is an N-terminal fusion to the target protein, a likely explanation for the absence of recovery is epitope masking/unavailability. Therefore, absence of a signal in the positive control was not considered to be a reason to fail the test antibody if all other results for that antibody fulfilled the passing criteria. For evaluating performance of mAb in IB application, a reagent was considered to pass if the signal was observed at the expected size in either the input (lanes 2 or 3) or IP lanes (lanes 5, 6 or 7), but not in the uninduced (lane1) and IgG-IP lanes (lane 4). The presence of a band at an unexpected size in either input or IP lanes was not considered negatively when evaluating the reagent. Such background signal in IB can likely be eliminated using standard techniques but must be evaluated by the end user of these reagents.

### Optimization and maintenance of hybridoma production for passing antibodies

Hybridoma cultures were expanded in medium containing DMEM (Cellgro,10-013-CM), 15% FBS (Cellgro, 35-016CV), P/S (Mediatech, MT 30-002-CI) and subcultured every 48h, while maintaining a minimum of 80% viability. Before freezing, cell lines were tested for Mycoplasma spp. contamination using PCR Mycoplasma Detection Kit (ABM, G238). If the result of the assay was negative, each cell line was expanded to a tissue culture flask of 175 cm^2^ and incubated for another 48h at 37°C/5%CO_2_. Cell lines with a density ranging from 5 – 8 × 10^5^ cells/mL, and with over 90% viability, were collected by centrifugation. Culture supernatant was stored at- 20°C for further testing, and cells were suspended in 10% DMSO (Sigma-Aldrich, D2650), 20% FBS (Cellgro, 35-011CV) and P/S (Mediatech, MT 30-002-CI) and frozen. Isotyping of each hybridoma cell line was done using Rapid Mouse Antibody Isotyping XpressCard (Antagen Pharmaceuticals, Inc., ISO-M8ac).Frozen hybridoma cell lines were shipped to the Developmental Studies Hybridoma Bank, University of Iowa.

### Antibody purification

Antibody from culture supernatant was affinity purified with Protein G beads according to manufacturer’s instructions (Sigma Aldrich, P7700). Concentration of the eluted IgG samples and buffer exchange were done by adding 3 column volumes of PBS (Corning Cellgro, 46-013-CM), 30% glycerol (Sigma Aldrich, G5516) and 0.02% sodium azide (Sigma Aldrich, S2002) to a Spin-X^®^ UF 6mL Centrifugal Concentrator, 50,000 MWCO Membrane (Corning, 431485).amples were reduced to a volume of 1mL, and antibody concentration was determined with aNanoDrop™ Lite Spectrophotometer (ThermoFisher Scientific).

### ChIP-seq

Exponentially growing cells from ENCODE Tier 1 and Tier 2 cell lines were crosslinked in 1% formaldehyde and harvested in batches of 20 million cells per replicate. Prepared nuclei were resuspended in RIPA buffer and chromatin shearing was performed on a Bioruptor Twin instrument (Diagenode). Antibody (5 μg/rxn) was coupled to M-280 Dynabeads (ThermoFisher Scientific; 11201) overnight prior to addition of sheared chromatin. A small aliquot of the sheared chromatin was reserved as an input control. Following incubation and wash steps, captured chromatin and input control were reverse crosslinked overnight at 65°C and recovered using the QIAquick PCR Purification System (Qiagen; 28104). ChIP-seq libraries were prepared and run on an Illumina HiSeq 2500 instrument. ChIP-seq biological replicates represent immunoprecipitations from two independent growths of the same cell line. Sequencing reads were aligned to the human hg19_Female genome using Bowtie2. The MACS2 algorithm was used to find significant peak signals above background (input control) and Homer was used to find enriched known and *de novo* motifs. A detailed protocol can be found at the ENCODE Data Portal: https://www.encodeproject.org/documents/6ecd8240-a351-479b-9de6-f09ca3702ac3/@@download/attachment/ChIP-seq_Protocol_v011014.pdf

### *Xist* labelling of HuProt

Cy5-UTP incorporated cRNA probes of *Xist* produced by T7-directed transcription was a kind gift from Eric Lander’s lab (MIT). Labeling and developing of the HuProt was performed as described previously^12^.

### ELISA, SPR and IP-MS experiments

Detailed protocols for these experiments can be accessed at https://proteincapture.org/protocols.

## Acknowledgements

This work was supported by NIH Common Fund Awards U54HG006434 (Johns Hopkins) and U01DC011485 (Rutgers). All ChIP-Seq data has been deposited in GEO

## Contributions

A.V.,M.M.,P.M.,Z.K.,L.X.,D.G.,S.L.,P.R.,S.H.,D.B.,R.S.,S.C.,Ho.Z.,F.P.,G.S.,J.N.,E.A.,L.A.,L.R.,L.L.,G.M.,J.R.,K.R.,R.A.,L.N.,K.M.,I.V.,Z.A.R.,C.R.,M.V.,J.M.,B.S.C.,S.Y.,S.G.K.,J.d.,M.S.,L.J.,A.T. and E.C. performed experimental work. K.Y., J.I. and Sa.K. designed algorithms and implemented software.

W.Y.Y.,S.A.,R.M.,J.D.B.,D.F.,G.W.,D.J.E.,J.S.,I.P.,H.Z. and S.B. contributed expertise and supervision. All authors contributed to manuscript preparation.

## Corresponding authors

Address correspondence to IP, DE, HZ and SB.

## References

1. Baker, M. Reproducibility crisis: Blame it on the antibodies. Nature 521, 274–276 (2015).

2. Bradbury, A. & Pluckthun, A. Reproducibility: Standardize antibodies used in research. Nature 518, 27–29 (2015).

3. Weller, M.G. Quality Issues of Research Antibodies. Anal Chem Insights 11, 21–27 (2016).

4. Pauly, D. & Hanack, K. How to avoid pitfalls in antibody use. F1000Res 4, 691 (2015).

5. Schonbrunn, A. Editorial: Antibody can get it right: confronting problems of antibody specificity and irreproducibility. Mol Endocrinol 28, 1403–1407 (2014).

6. Bordeaux, J. et al. Antibody validation. Biotechniques 48, 197–209 (2010).

7. Saper, C.B. & Sawchenko, P.E. Magic peptides, magic antibodies: guidelines for appropriate controls for immunohistochemistry. J Comp Neurol 465, 161–163 (2003).

8. Na, H. et al. A high-throughput pipeline for the production of synthetic antibodies for analysis of ribonucleoprotein complexes. RNA 22, 636–655 (2016).

9. Hornsby, M. et al. A High Throughput Platform for Recombinant Antibodies to Folded Proteins. Mol Cell Proteomics 14, 2833–2847 (2015).

10. Marcon, E. et al. Assessment of a method to characterize antibody selectivity and specificity for use in immunoprecipitation. Nat Methods 12, 725–731 (2015).

11. Rhodes, K.J. & Trimmer, J.S. Antibodies as valuable neuroscience research tools versus reagents of mass distraction. J Neurosci 26, 8017–8020 (2006).

12. Uhlen, M. et al. Proteomics. Tissue-based map of the human proteome. Science 347, 1260419 (2015).

13. Blackshaw, S. et al. The NIH Protein Capture Reagents Program (PCRP): a standardized protein affinity reagent toolbox. Nat Methods 13, 805–806 (2016).

14. Hu, C.J. et al. Identification of new autoantigens for primary biliary cirrhosis using human proteome microarrays. Mol Cell Proteomics 11, 669–680 (2012).

15. McHugh, C.A. et al. The Xist lncRNA interacts directly with SHARP to silence transcription through HDAC3. Nature 521, 232–236 (2015).

16. Chu, C. et al. Systematic discovery of Xist RNA binding proteins. Cell 161, 404–416 (2015).

17. Vaquerizas, J.M., Kummerfeld, S.K., Teichmann, S.A. & Luscombe, N.M. A census of human transcription factors: function, expression and evolution. Nat Rev Genet 10, 252–263 (2009).

18. Greenspan, N.S. Cohen’s Conjecture, Howard’s Hypothesis, and Ptashne’s Ptruth: an exploration of the relationship between affinity and specificity. Trends Immunol 31, 138–143 (2010).24

19. Steward, M.W. & Lew, A.M. The importance of antibody affinity in the performance of immunoassays for antibody. J Immunol Methods 78, 173–190 (1985).

20. Abdiche, Y., Malashock, D., Pinkerton, A. & Pons, J. Determining kinetics and affinities of protein interactions using a parallel real-time label-free biosensor, the Octet. Anal Biochem 377, 209–217 (2008).

21. Mita, P. et al. Fluorescence ImmunoPrecipitation (FLIP): a Novel Assay for High-Throughput IP. Biol Proced Online 18, 16 (2016).

22. Jeong, J.S. et al. Rapid identification of monospecific monoclonal antibodies using a human proteome microarray. Mol Cell Proteomics 11, O111 016253 (2012).

23. de Melo, J. et al. Lhx2 Is an Essential Factor for Retinal Gliogenesis and Notch Signaling. J Neurosci 36, 2391–2405 (2016).

24. Harlow E., Lane D. Using Antibodies: A Laboratory Manual, Cold Spring Harbor Laboratory, Cold Spring Harbor, NY (1998).

25. Uhlen, M. et al. A proposal for validation of antibodies. Nat Methods 13, 823–827 (2016).

26. Roncador, G. et al. The European antibody network’s practical guide to finding and validating suitable antibodies for research. MAbs 8, 27–36 (2016).

27. GBSI Workshop Report: Antibody Validation: Strategies, Policies, and Practices at https://www.gbsi.org/gbsi-content/uploads/2016/12/Workshop-Report12-15-2016.pdf

## References for Online Methods

1. Vaquerizas, J.M., Kummerfeld, S.K., Teichmann, S.A. & Luscombe, N.M. A census of human transcription factors: function, expression and evolution. Nat Rev Genet 10, 252–263 (2009).

2. Hu, S. et al. Profiling the human protein-DNA interactome reveals ERK2 as a transcriptional repressor of interferon signaling. Cell 139, 610–622 (2009).

3. Zhu, H. et al. Global analysis of protein activities using proteome chips. Science 293, 2101–2105 (2001).

4. Harlow E., Lane D. Using Antibodies: A Laboratory Manual, Cold Spring Harbor Laboratory, Cold Spring Harbor, NY (1998).

5. Jeong, J.S. et al. Rapid identification of monospecific monoclonal antibodies using a human proteome microarray. Mol Cell Proteomics 11, O111 016253 (2012).

6. Taylor, M.S. et al. Affinity proteomics reveals human host factors implicated in discrete stages of LINE-1 retrotransposition. Cell 155, 1034–1048 (2013).

7. Dai, L., Taylor, M.S., O’Donnell, K.A. & Boeke, J.D. Poly(A) binding protein C1 is essential for efficient L1 retrotransposition and affects L1 RNP formation. Mol Cell Biol 32, 4323–4336 (2012).

8. Mita, P. et al. Fluorescence ImmunoPrecipitation (FLIP): a Novel Assay for High-Throughput IP. Biol Proced Online 18, 16 (2016).

9. Longo, P.A., Kavran, J.M., Kim, M.S. & Leahy, D.J. Transient mammalian cell transfection with polyethylenimine (PEI). Methods Enzymol 529, 227–240 (2013).

10. de Melo, J. et al. Injury-independent induction of reactive gliosis in retina by loss of function of the LIM homeodomain transcription factor Lhx2. Proc Natl Acad Sci U S A 109, 4657–4662 (2012).

11. Lee, D.A. et al. Tanycytes of the hypothalamic median eminence form a diet-responsive neurogenic niche. Nat Neurosci 15, 700–702 (2012).

12. Rapicavoli, N.A., Poth, E.M., Zhu, H. & Blackshaw, S. The long noncoding RNA Six3OS acts in trans to regulate retinal development by modulating Six3 activity. Neural Dev 6, 32 (2011).

